# Notch, β-catenin and MAPK signaling segregate endoderm and mesoderm in the diploblast *Nematostella vectensis*

**DOI:** 10.1101/2024.10.29.620801

**Authors:** Emmanuel Haillot, Tatiana Lebedeva, Julia Steger, Grigory Genikhovich, Juan D. Montenegro, Alison G. Cole, Ulrich Technau

## Abstract

Cnidaria are typically considered diploblastic (i.e. consisting of two germ layers) in contrast to their triploblastic sister clade, the Bilateria. However, a recent study suggested that sea anemones and other cnidarians have three segregated germ layer identities, corresponding to the bilaterian germ layers^1^. Here, we investigated, how the three germ layer identities are specified during early development of the sea anemone *Nematostella vectensis*. First, the mesodermal territory is specified at the animal pole at 6 hours postfertilization, followed by the specification of a ring of endodermal territory between mesoderm and ectoderm. We then assessed the role of β-catenin, MAPK and Notch signaling pathways during mesoderm and endoderm formation. Our results show that the mesoderm is initiated by MAPK signaling and simultaneously restricted to the future oral side by mutually exclusive nuclear β-catenin signaling. The mesodermal cells then express the Delta ligand, while the ectodermal cells express the Notch receptor. Inhibition of Notch signaling as well as ectopic expression of the Notch intracellular domain showed that endodermal tissue identity is induced by Notch signaling at the boundary between mesoderm and ectoderm. We propose a new model that outlines the different steps leading to the segregation of mesoderm and endoderm identities in *Nematostella*, confirming the presence of 3 distinct germ layer identities in this cnidarian. Notably, the observed crosstalk of MAPK, β-catenin and Notch signaling in the specification of three germ layers in *Nematostella* is highly reminiscent to early stage gastrulae of sea urchins suggesting that triploblasty may predate the split of cnidarians and bilaterians.

## Introduction

All bilaterians are composed of three embryonic germ layers, ectoderm, endoderm and mesoderm, which give rise to all tissues in the adult organism. In contrast, Cnidaria, the sister group of Bilateria, consist of only two cell layers. Traditionally, it was assumed that the two cell layers of cnidarians are homologous to bilaterian endoderm and ectoderm, and the mesoderm arose at the base of Bilateria. To trace the evolutionary origin of mesoderm, conserved transcription factors involved in mesoderm formation and differentiation in Bilateria were cloned from various cnidarians ^2–6^. Most of these genes are expressed in the inner cell layer, which led to the suggestion that the inner layer constitutes a mesendoderm, an evolutionary precursor of mesoderm and endoderm. However, expression analyses of many more conserved endodermal and mesodermal markers throughout development of the sea anemone *Nematostella vectensis* challenged this view and showed that inner layer has a molecular profile reminiscent of the bilaterian mesoderm, and gives rise to muscles, gonads and nutrient storage, typical derivatives of mesoderm in Bilateria. Conversely, the pharyngeal ectoderm and its derivatives, the septal filaments at the distal tip of the mesenteries, expresses many conserved endodermal markers and differentiates into digestive gland cells and insulin-secreting cells, characteristic of endoderm derivatives in Bilateria ^1^. These findings have challenged the presumed homology of cnidarian inner and outer cell layers with ectoderm and endoderm.

The specification of endoderm and mesoderm is the first cell fate decision in all embryos and hence a key step in animal development. In ambulacrarian deuterostomes, such as echinoderms and hemichordates, in urochordates and at least in some spiralian protostomes such as nemertines, endomesoderm formation is governed by β-catenin signaling ^7–10^. Notch signaling was shown to be crucial in the segregation of mesendoderm into endoderm and mesoderm in echinoderms ^9,11,12^ and mitogen-activated protein kinase (MAPK) signaling is also involved in specification of the mesoderm in Ambulacraria, ecdysozoans and spiralians, suggesting an ancestral role in the common bilaterian ancestor ^13^ (Figure 7). Thus, the regulatory relationships between these pathways may vary, however, their involvement in the bilaterian and cnidarian germ layer specification is a recurrent theme.

## Materials and Methods

### Animal culture and spawning

Adult polyp of *Nematostella vectensis* were cultivated in 16‰ Red Sea Salt artificial sea water, (*Nematostella* medium=NM) at 18°C in the dark, according to the protocol described ^17^. Animals were fed 5x per week with freshly hatched *Artemia* nauplii. Spawning in mature polyps was induced by exposure to light over 9 hours at 25°C. After fertilization, the jelly surrounding the eggs was removed with a 3% L-cystein/NM solution during 30 minutes, and the emrbyos were washed 5-6 times in NM and raised until required stage at 21°C as previously described ^3,17^.

### Generation of single cell transcriptomic data

Embryos were collected in Eppendorf tubes pre-coated with 0.1% PBST and washed with NM (1x for 12hpf, 2x for 8 and 10hpf). For cell dissociation at 8 and 10hpf, embryos were washed 3 times with 200mL NM containing 2% L-cystein (Millipore, #52-90-4), pH 7.5. Then embryos were kept at 4°C on ice and mixed every 3 minutes for 15 minutes. Cell suspension was centrifuged at 400 rcf for 30 seconds and L-Cystein solution was removed. Then cells were resuspended in 0.5% BSA/1xPBS by pipetting. This step was repeated 2 times. For cell dissociation at 12hpf, embryos were washed with NM and first incubated for 1 minute in 5x TrypLE (diluted in NM). Dissociation was achieved by gentle pipetting for 25 minutes and stopped by adding an equal volume of 1% BSA/1xPBS. Then, the cell supsension was centrifuged (200 rcf, 4 min), washed with 0.5% BSA/1xPBS, and passed through a Flowmi® filter (40µm). After dissociation and washing, concentration and viability of all cell suspensions was determined by staining cells with ViaStain AOPI.

### Analysis of single cell transcriptomic data

Reads were aligned to the *Nematostella vectensis* genome ^18^ using Cell Ranger v 7.1.0 ^19^ using standard parameters. The filtered count matrix was upload to Seurat v5 ^20–22^. Each library was filtered and normalized separately removing cells with low read counts and low feature counts. Cell cycle markers were also regressed out using the Cell Cycle Scoring method and new PCA and UMAP values were calculated. A detailed script can be found in our github (https://github.com/technau/NemVecEndoderm) for replication of our results.

### Signaling pathway inhibitor treatments

All stock solutions of inhibitor drugs were prepared in DMSO. Before first cell division, dejellied zygotes were transferred in a Petri dish with the appropriate concentration of inhibitor drugs. Zygotes were treated with a selective inhibitor of MEK (U0126, Sigma, #19-147) at 20μM to inhibit the MAPkinase signaling, a selective inhibitor of GSK3ß (1-azakenpaullone (AZ), Sigma) at 10μM to stabilize β-catenin, and with a selective inhibitor of γ-secretase (LY411575, MedChemExpress, #HY-50752) at 10-40μM to inhibit the Notch signaling.

### Microinjection of morpholinos, plasmid and shRNA

For knockdown experiment, a previously published antisense translation blocking Nv β-catenin morpholino (MO) (Gene Tools Inc., USA) was microinjected in zygotes at a concentration of 500μM with 0,125mg/ml of fluorescent Dextran-AlexaFluor488 (5’UTR Nv β-catenin Mo: 5’ TTCTTCGACTTTAAATCCAACTTCA 3’) ^15^. As a control, a previously used standard morpholino known to have no effect on the development was injected at 500μM (standard MO 5’ GATGTGCCTAGGGTACAACAACAAT 3’) as previously described ^23,24^. For mosaic overexpression of the Notch intracellular domain (NICD), injection of TBP::NICD-mCherry transgenic vector plasmid in zygotes was performed as previously described ^25^. To generate this vector, the cDNA fragment coding for the NICD was amplified with insertion of an ATG at the beginning. This sequence was fused to mCherry sequence with the 4xGly-1xSer linker (5’GGTGGTGGTGGTAGT‘3) and inserted in the pJET1.2 plasmid downstream a T7 RNA polymerase site by using NEBuilder HIFI DNA Assembly Cloning Kit (NEB; #E5520S). Gene knockdown mediated by shRNA was realized as described previously ^26^. ShRNA was used at 500ng/μL. shRNA against mOrange was used as a control.

### Antibody and phalloidin staining

For immunostaining, embryos were fixed for 1 h with 4% PFA/PBST (4% paraformaldehyde dissolved in 1xPBS, 0.4% Tween 20) at room temperature without agitation and washed 4 times with PBST. Then, embryos were incubated in blocking solution (5% sheep serum, 1% bovine serum albumin (BSA) in PBST for 1h at room temperature (RT). Embryos were incubated overnight at 4°C in anti phospho-ERK (Cell Signaling Technology; #4370S, 1/500) or anti NICD (Cell Signaling Technology; #4147T; 1/500) diluted in the blocking solution. After washing 7 times for 10 minutes with PBST, embryos were incubated in the blocking solution with anti-rabbit AP (Invitrogen; #31346, 1/7000) overnight at 4°C. After washing 10 times for 10 minutes, staining was revealed with NBT/BCIP in AP buffer. For phalloidin staining, after fixation in 4% PFA and washing with PBST, embryos were incubated in 1ml acetone on ice for 5 minutes. Specimen were washed 5 times with PBST, then incubated with 100μL PBST containing 4 U/mL Phalloidin Alexa Fluor 488 (Thermo Fisher Scientific) and 5 µg/mL DAPI in the dark for 1h at RT. Then, embryos were washed 7 times for 10 minutes with PBST followed by infiltration with antifade mounting medium (Vectashield). Imaging was performed with a Nikon Eclipse 800 or Leica SP8 CLSM.

### Western Blot

To generate lysates, embryos were incubated in 40μl of cell extraction buffer (FNN0011, ThermoFisher) supplemented with protease inhibitor (Roche). Samples were centrifuged at 16000 g for 10 minutes at 4°C and supernatants were resuspended in 10μl loading dye. After gel electrophoresis and blotting, the membranes were blocked in 5% skimmed milk powder in PTw (1xPBS, 0.1% Tween). Antibodies against ERK (#4695, Cell Signaling), phospho-ERK (#4370, Cell Signaling) and β-actin (#4970S, Cell Signaling) were diluted at 1:10000 in blocking solution and incubated with Nitrocellulose membrane overnight at 4°C. After washing with PTw (1x PBS, 0.1% Tween20), membranes were incubated with1:100000 anti-rabbit IgG conjugated to horseradish peroxidase (#A0545, Sigma-Aldrich), washed with PTw, and peroxidase activity associated with proteins of interest was detected with SuperSignal West Femto Maximum Sensitivity Substrate (#34094, Thermo Fisher).

### Cloning and *in situ* hybridization

To generate probes for in situ hybridization, transcripts of interests were amplified from cDNA using primers listed in Table 1. PCR products were cloned in pGEM-T vector (Promega, #A3600). FITC or DIG-labeled RNA probes were generated by in vitro transcription with SP6 or T7 polymerase. In situ hybridization was carried out as previously described ^23^ with some modifications.

After incubation with anti-Digoxigenin AP (1/4000), embryos were washed 10 times 10 minutes with TBST (0.4% Tween). Background was removed by washing embryos 6 times with TBST (0.4% Tween) and kept overnight at 4°C. Two-color fluorescent in situ hybridization was performed similarly to single in situ hybridization according to the protocol described with some following differences ^27^. Fluorescent staining was revealed with the TSA Plus Fluorescein and TSA Plus Cy3 detection Kits (AKOYA Biosciences). After staining, embryos were infiltrated with Vectashield (VectorLabs) and imaged using Leica SP8 CLSM.

## Results

### Mesoderm and ectoderm segregate before the endoderm

*Nematostella* eggs contain maternal mRNAs of genes, whose zygotic expression later marks aboral ectoderm and becomes gradually cleared first from the mesodermal domain and later from the endoderm, oral and midboby ectoderm, as they arise ^24^. Previous studies identified a number of specific marker genes for endoderm or mesoderm ^1,2,4,28^. To determine the precise timing of mesoderm and endoderm initiation during development and to define more clearly the key steps responsible for their emergence, we generated single cell RNA libraries of 8, 10 and 12hpf. Additionally, single cell RNAseq data previously generated by our lab at 18, and 24 hpf ^29,30^ were also used for this analysis. After cell clustering, we were able to distinguish the cell populations with ectodermal, mesodermal and endodermal identities. Notably, at 8hpf, only clusters of mesodermal and ectodermal cells are clearly identifiable (Figure S1). Our data show sequential activation of mesodermal genes (Figure 1B). The first expression of zygotic genes encoding TFs such as *tbx19-like*, *gsc2-like*, *duxABC1* and *duxABC2* starts around 6hpf. By 8hpf, *pitx1-like*, *fgfa1*, and *erg* are turned on in addition (Figures 1A and 1D). This is followed by the expression of *nkx2.2B* detectable at 10hpf and by the expression of new mesodermal genes such as *zicA*, *snailA*, *mitf-like*, *runx*, *six4*, and *smad1/5* by 12hpf. At 14hpf *hand2, zc3h12-like*, *nkx2.2D*, *otxB*, and at 16hpf *isx-like 2*, *hmx2* and *otxC* become detectable, shortly before the onset of gastrulation (Figure 1B and 1D). The ectodermal cells express genes encoding homeobox transcription factors such as *koza-like1/2*, or a *bHLH TF-like* in a pattern complementary to the mesodermal gene expression. Initiation of the endodermal germ layer identity can be clearly detected at 10 hpf (*brachyury*) and 12hpf (*foxA*, *wnt1* and *wnt3*) in the cells surrounding the mesoderm (Figure 1C). From 12hpf until 20hpf, endodermal markers, *foxA*, *wnt1* and *wnt3* are expressed in a one cell diameter-wide ring surrounding the mesoderm, and it is only at the onset of gastrulation that the endodermal domain will expand (Figure 1C). As previously observed ^23^, the very early mesodermal cells also transiently and very weakly express *wnt1, wnt3 and wntA*, which are future markers of endoderm and later associated with maintaining the endodermal fate. In line with this faint expression, we could not detect any Wnt signaling target genes characteristic of endodermal fate, such as *brachyury and foxA* in the future mesoderm (Figure 1C). Regardless, the early weak expression of *wnt1, wnt3, wntA* is rapidly cleared from the mesodermal plate and forms a ring in the future endoderm by 12-14 hpf (Figure 1C). This raises the possibility that the endoderm could be specified by the interaction between mesoderm and ectoderm. Together, this in situ hybridization screen shows that there is an approximately 4 hours delay between the onset of the mesodermal marker gene expression at 6-8 hpf and the start of the endodermal marker gene expression at 10-12 hpf (Figure 1D).

**Figure 1.**
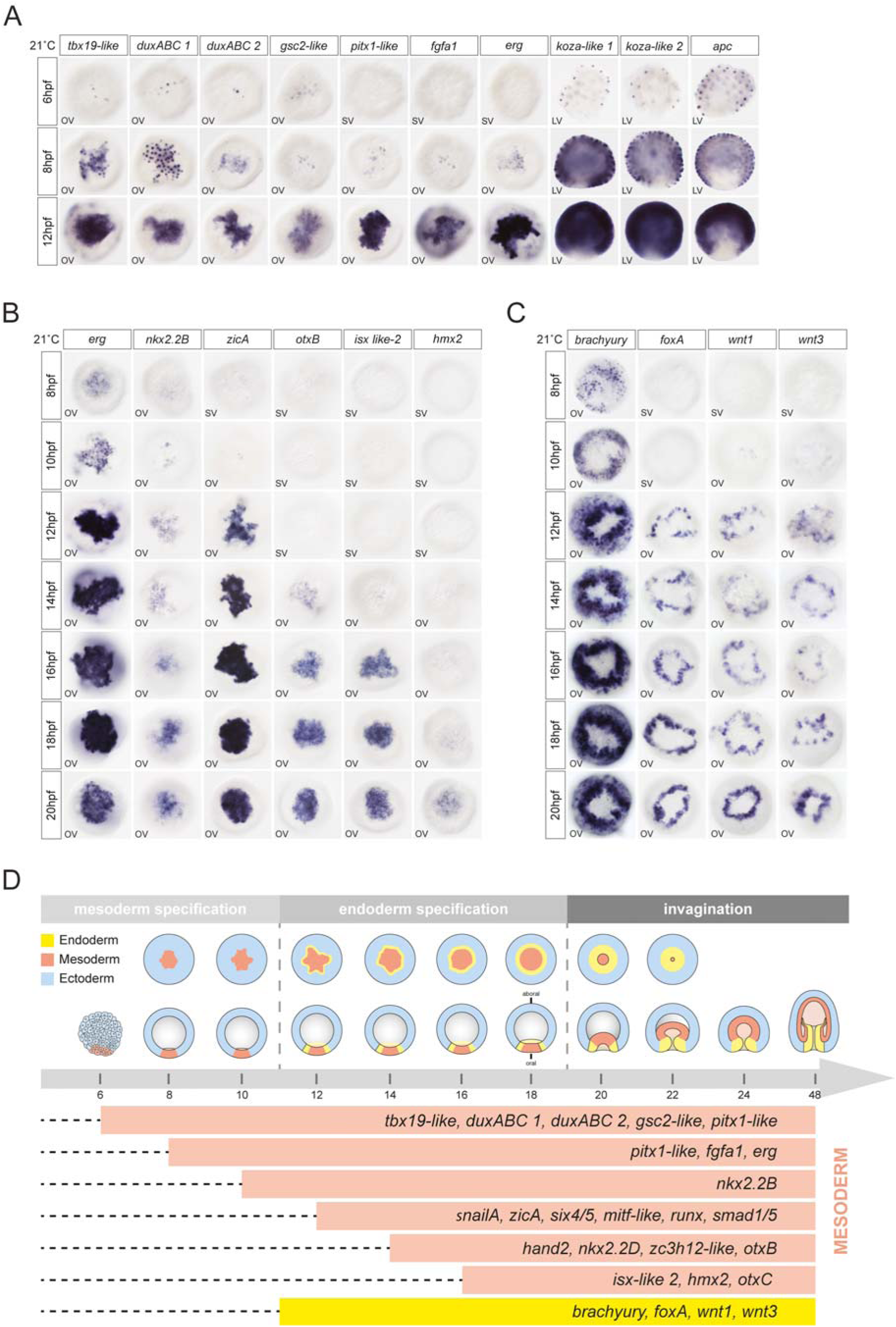
Expression profiles of ectodermal, mesodermal, and endodermal genes at early embryonic stages in *Nematostella*. (A) Spatio-temporal expression patterns of mesodermal and ectodermal markers at 6, 8 and 12hpf revealed by in situ hybridization. OV; oral view, SV; surface view. (B) Spatio-temporal expression patterns of mesodermal TFs at 8, 10, 12, 14, 14, 16, 18, 20, and 26hpf. OV; oral view, SV; surface view. (C) Spatial and temporal expression profiles of the endodermal TFs during early development. OV; oral view, SV; surface view. (D) Summary of temporal succession of mesodermal and endodermal gene expression profiles during early development of *Nematostella*.

### β-catenin signaling promotes ectoderm and restricts mesodermal identity at the future oral pole

The delay in the onset of the endodermal marker gene expression in comparison to the mesodermal markers suggested a difference in their regulation and prompted us to investigate Wnt/β-catenin and MAP kinase signaling pathways, which were previously implicated in the development of the mesoderm in *Nematostella*. Some studies ^14,28,31^ suggested that β-catenin signaling promotes the specification of mesoderm, however, others presented experimental evidence showing that it antagonizes mesoderm formation ^15,16^. Inactivation of MAPK signaling also perturbed mesoderm development and gastrulation movements ^30^, however, the role of MAPK in relation to Wnt signaling in germ layer formation was not addressed in detail.

To clarify the role of the Wnt/β-catenin pathway in the process of mesoderm and endoderm formation, we analyzed the expression of earliest mesodermal genes by in situ hybridization upon down- or up-regulation of the Wnt/β-catenin pathway (Figures 2A and 2B). We analysed the effects of these treatments at 8 hpf, when only ectoderm and mesoderm is specified, and at 20 hpf early gastrula, when all three germ layers are present and mesoderm starts to invaginate during normal development. While untreated embryos display localized expression of *fgfa1, tbx19-like, duxABC* and *pitx1-like* in the future mesoderm at 8 hpf, knockdown of β-catenin by injection of a translation-inhibiting morpholino results in an ectopic expression of these mesodermal genes throughout the embryo, while ectodermal markers such as koza1-like and APC are downregulated (Figure 2C). Conversely, ectopic upregulation of the Wnt/β-catenin pathway by treatment with the GSK3β inhibitor Azakenpaullone (AZ) abolishes mesodermal marker expression and leads to ubiquitous expression of ectodermal markers compared to control (Figure 2C). Hence, in line with the earlier reports ^15,16^, we conclude that at this early stage β-catenin represses the mesodermal program and promotes the ectodermal program, restricting the mesoderm to the domain at the animal pole, where nuclear β-catenin is not expressed. The “anti-mesodermal” effect of β-catenin persists to later stages. At 20 hpf, mesodermal genes extend throughout the embryo upon β-catenin knockdown and disappear after upregulation of the β-catenin pathway by AZ treatment (Figure 2D). The striking difference between the effects of β-catenin pathway manipulation at 8 hpf and 20 hpf is its effect on the emerging endoderm. Expression of the endodermal markers *foxA* and *brachyury* is abolished upon β-catenin knockdown and extends throughout the whole embryo in azakenpaullone treated embryos, indicating that endodermal identity depends on β-catenin signaling (Figure 2E). Together, we confirm that high levels of β-catenin promote endodermal fate in the oral ectoderm at the expense of aboral ectoderm, which requires lower levels of β-catenin ^15,24,33^.

**Figure 2.**
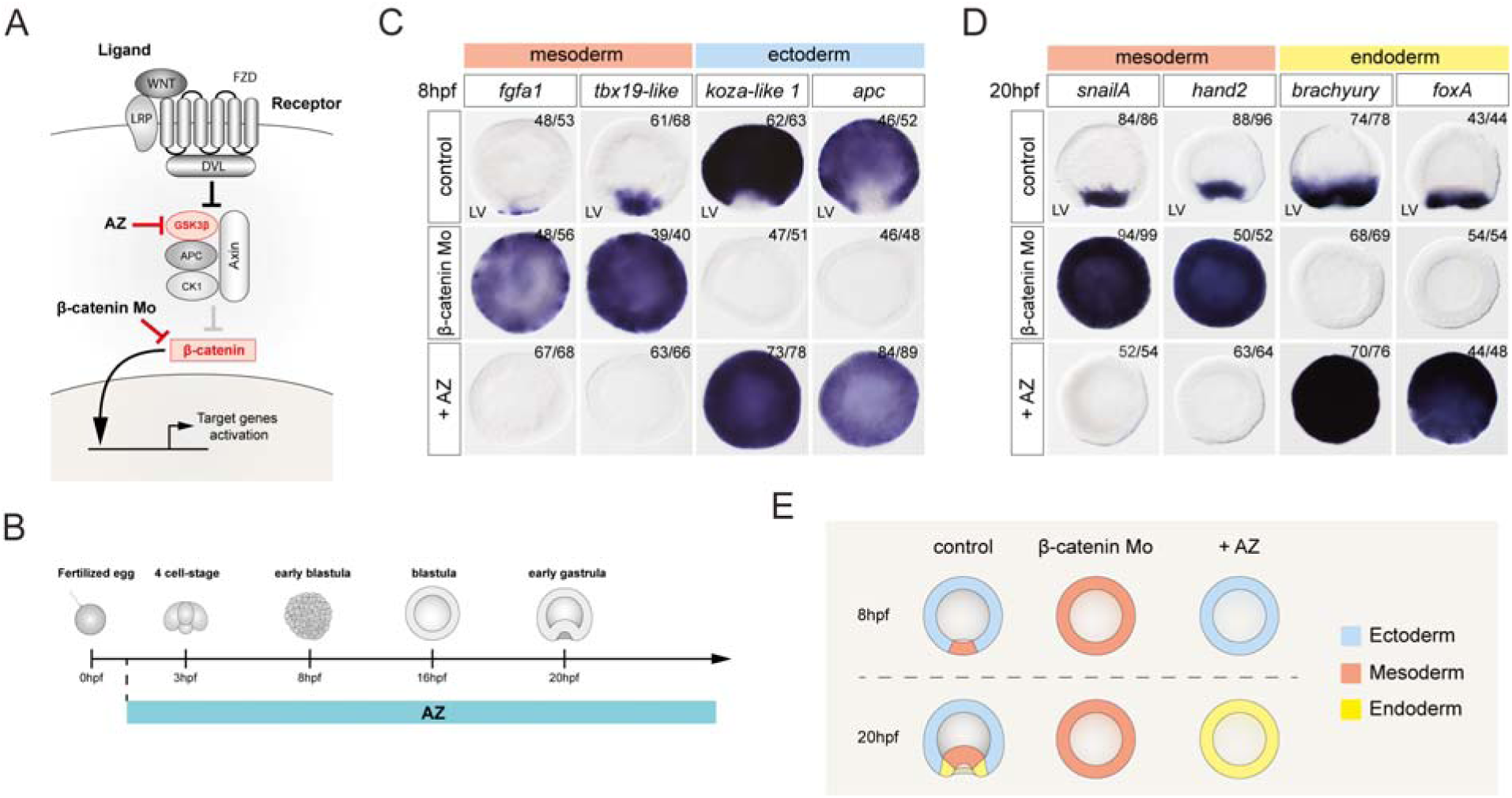
β-catenin signaling restricts mesoderm fate in oral pole at early stage. (A) Scheme of the Wnt/ β-catenin signaling pathway and the role of the GSK3β inhibitor Azakenpaullone (AZ) and of β-catenin morpholino. (B) Scheme of treatment with AZ during the embryonic development. (C) In 8hpf embryos, mesodermal markers *fgfa1* and *tbx19-like* become ubiquitously expressed upon beta-catenin knockdown and are abolished upon AZ-mediated beta-catenin stabilization. Ectodermal markers koza-like1 and apc demonstrate an opposite effect. LV; lateral view. (D) Expression of mesodermal (*snailA*, *hand2*) and endodermal (*brachyury*, *foxA*) genes in AZ treated embryos or β-catenin morpholino embryos at 20hpf. LV; lateral view. (E) Scheme describing the fate map changes induced after up and downregulation of β-catenin at 8hpf and 20hpf.

### MAPK signaling is essential for mesoderm induction and invagination

Based on previous studies, the MAP kinase pathway emerges as a potential candidate in the process of mesoderm formation ^32,34^, however, its function and how it interacts with the β- catenin pathway was unclear. To elucidate the role of the MAP kinase pathway (Fig. 3A), we identified the areas of active MAPK signaling during the mesoderm formation by immunolabelling the activated form of ERK (pERK) and by in situ hybridization against the MAPK target gene *erg*. We were able to detect pERK from 6-8 hpf on in the nuclei of the mesodermal cells, promptly followed by the expression of the MAPK target gene *erg* (Figure 3B). Western blot analyses confirmed that the MEK inhibitor U0126 can inhibit MAPK signaling and abolish the phosphorylation of ERK within 30 minutes of treatment (Figures 3C and 3D). Embryos treated with U0126 upon fertilization showed a loss of expression of mesodermal markers *erg*, *tbx19-like* and *fgfa1* (Figure 3D). This observation at early blastula suggests that the MAP kinase pathway is required for the mesoderm specification. Prolonged treatment with U0126 until the onset of gastrulation (20 hpf) resulted in the abolishment of most of the mesodermal genes, although *erg*, *six4/5* and *zicA* maintained their expression in a smaller oral domain (Figures 4A and S4A). By comparison, the expression of endodermal markers, such as *foxA* and *wnt3* is still present in U0126 treated embryos. Moreover, we observe ectopic expression of *brachyury* and *foxA* in the mesodermal territory suggesting that these genes are normally suppressed in the mesoderm by MAPK signaling (Figure 4A, see also ^32^). Similar results were obtained after downregulation of the ERG expression, one of the key transcription factors downstream of the MAPK signaling ^34^ (Figure S4B).

**Figure 3.**
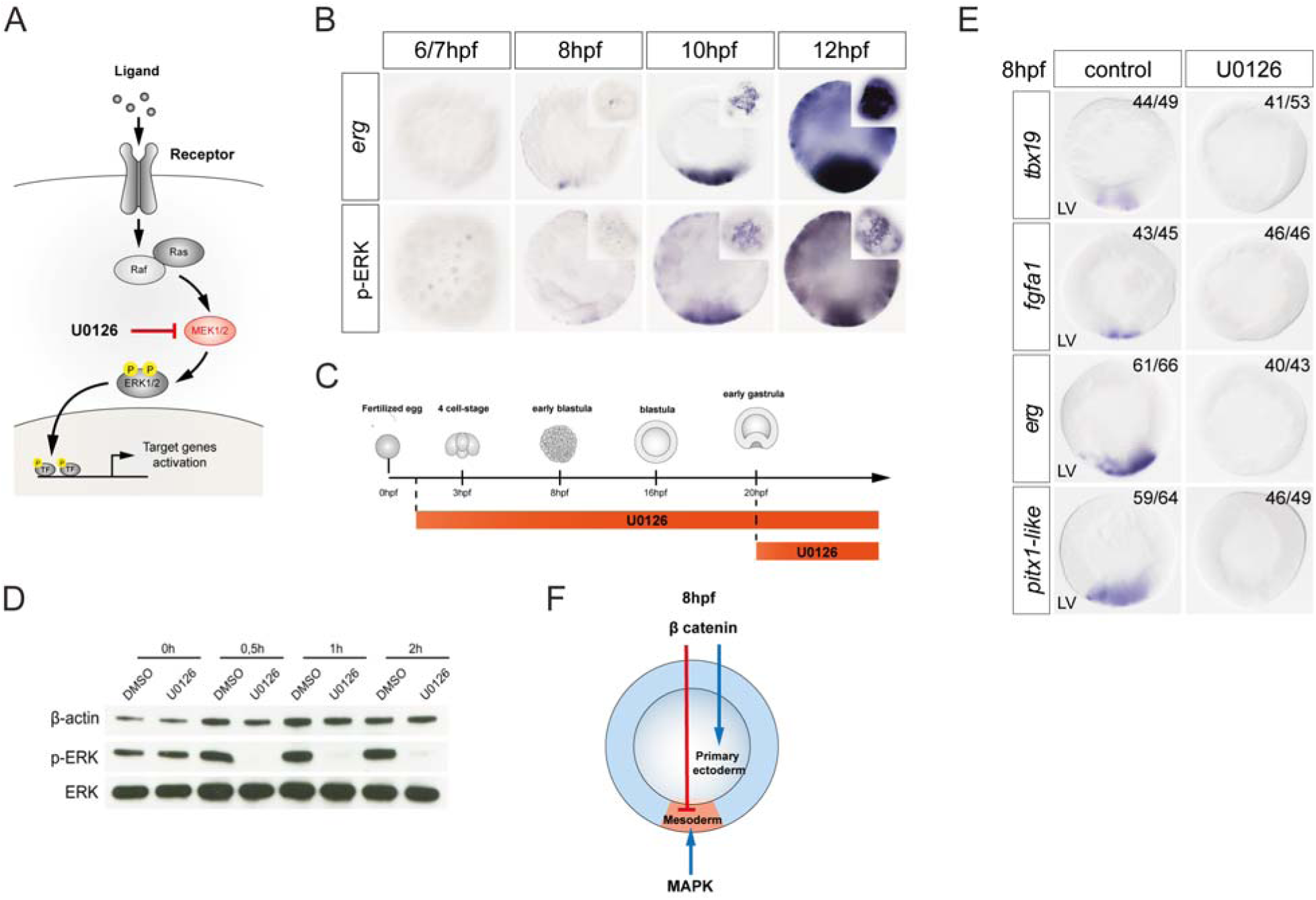
MAP kinase signaling is essential for mesoderm induction. (A) Scheme of the RTK/MAPK signaling pathway and the role of the MEK inhibitor U0126. (B) Spatial and temporal expression profile of *erg* revealed by in situ hybridization in comparison to phospho-ERK immunostaining at 6, 8, 10, 12hpf. (C) scheme of treatments with U0126. (D) Western blot analysis of lysates of DMSO- or U0126-treated 20 hpf embryos shows that U0126 efficiently blocks ERK phosphorylation after a 30 minute treatment. (E) Mesodermal (*tbx19-like, fgfa1, erg, pitx1-like*) gene expression is suppressed by the U0126 mediated inhibition of MAPK signaling at 8hpf. LV; lateral view. (F) Schematic summary of the role of β-catenin and MAPK signaling in the specification of mesoderm.

**Figure 4.**
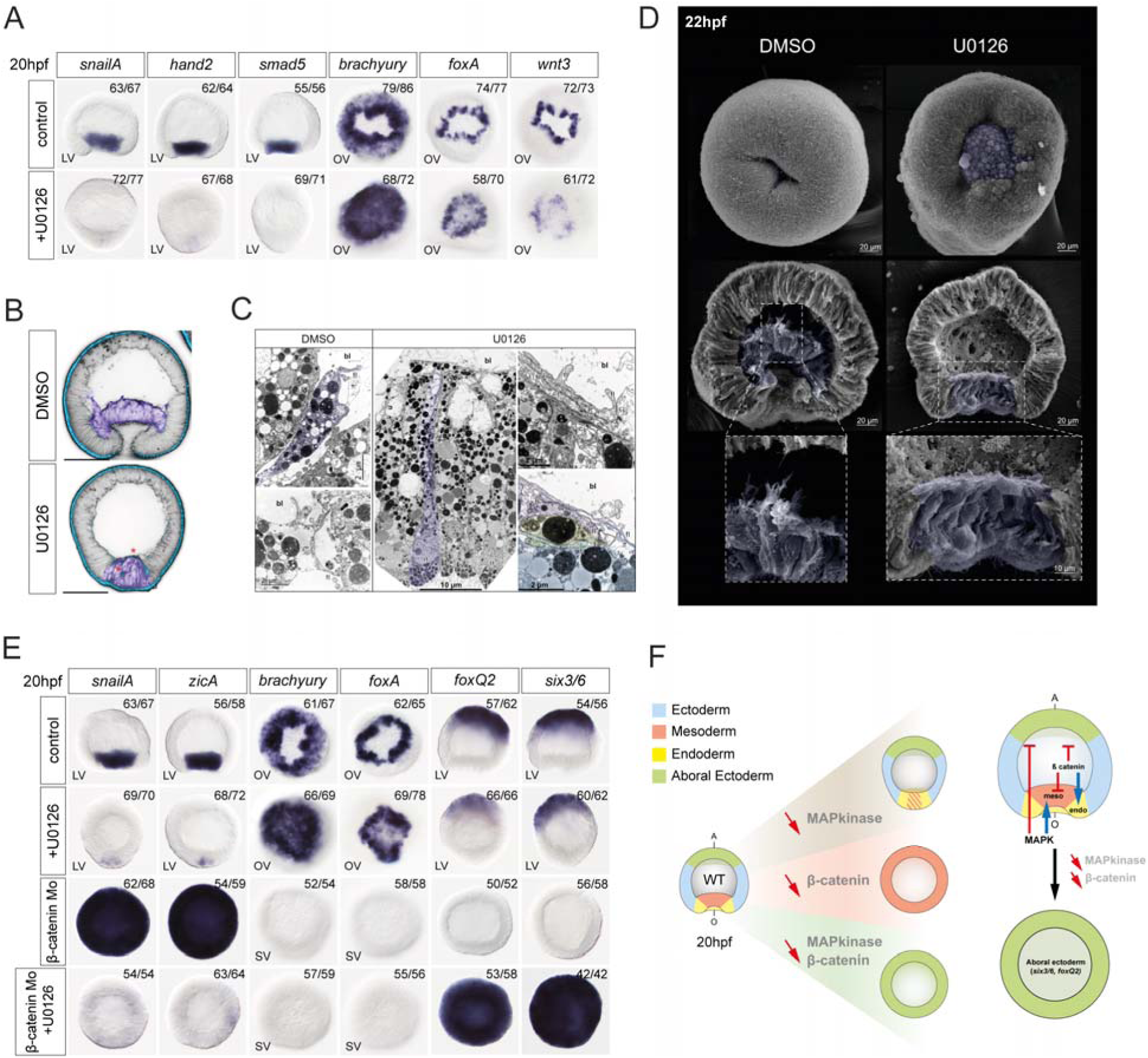
MAP kinase signaling is essential for mesoderm initiation and specification but also to inhibit aboral ectoderm. (A) At 20hpf, MAPK signaling inhibition with U0126 abolished expression of mesodermal markers and expands expression of endodermal markers (*brachyury*, *foxA*) into the mesodermal domain. LV; lateral view, OV; oral view. (B) Ectodermal cadherin cdh3 (blue) does not disappear from the mesodermal cells upon MAPK signaling inhibition. Cell outlines are shown by fibrillar actin staining (black). Mesodermal cells are highlighted in purple. Scale bars 50 µm. (C) TEM comparison of the U0126-treated and control DMSO-treated embryos at early gastrula stage shows that in U0126, mesodermal cells do not become bottle-shaped, and their filopodia appear to stick together and form numerous contacts rather than extend into the blastocoel. bl; blastocoel, fl; filopodia, cj; cell junction, n; nucleus. (D) SEM images of the U0126 and DMSO treated embryos at mid-gastrula stage. Note the lack of mesoderm invagination into the blastocoel and the absence of extending filipodia in U0126 treated embryos. (E) At 20hpf, MAPK and β-catenin signaling inhibition abolished mesodermal and endodermal markers but expands ubiquitously expression of ectodermal markers (*foxQ2*, *six3/6*). LV; lateral view, OV; oral view, SV; surface view. (F) Schematic summary of single or combined knockdown/inhibition of β-catenin and MAPK signaling on aboral ectoderm marker genes (*foxQ2*, *six3/6*).

Although changes in gene expression show that inhibition of the MAPK pathway by U0126 treatment affects the specification of mesodermal identity at the molecular level, the mesoderm nevertheless becomes morphologically distinct at the time of gastrulation. However, gastrulation is arrested shortly after the onset (Figure 4B-4D). During normal gastrulation, mesodermal cells become loosely packed, acquire bottle-cell morphology, and form filopodia reaching to the basal surfaces of the ectodermal cells as the mesoderm and ectoderm extend their contact in a zipping motion ^35,36^. Ultrastructural analyses of U0126 treated embryos show that the mesodermal cells start showing signs of the incomplete EMT (e.g. onset of the cell shape change, nuclear migration) but fail to disassemble their basal contacts and extend filopodia to the basal surfaces of the ectodermal cells (Figures 4B-4D). This suggests that cell- cell adhesion is affected by the inhibition of the MAPK pathway. Normal gastrulation in *Nematostella* is associated with a switch of Cadherin3 to Cadherin1 in the mesoderm ^37^. Embryos with MAPK pathway inhibited through treatment with U0126 or knockdown of *ERG* maintain Cadherin3 expression in the mesoderm, which may explain the retention of basal contacts between the mesodermal cells, the lack filopodia extension and of the migration of the mesodermal cells along the blastocoel roof (Figures 4B and S4C). Thus, MAPK signaling is essential to the specification of the mesoderm and subsequent morphogenetic processes required for gastrulation.

### MAPK and **β**-catenin signaling suppress the default aboral ectodermal identity of the embryo

Upon MAPK signaling suppression, Wnt/β-catenin targets *brachyury* and *foxA* expand into the mesodermal domain, suggesting that endodermal fate is suppressed in the area of active MAPK signaling. However, we wondered, whether endoderm had the same suppressive effect on the mesodermal fate. To this end, we injected zygotes with β-catenin morpholino, which normally should activate the mesodermal program throughout the entire embryo and downregulate endodermal genes, but in parallel we inhibited the MAP kinase pathway using U0126 and analyzed the expression of marker genes for the endoderm, the mesoderm and the aboral ectoderm (Figure 4E). In this condition of simultaneous β-catenin knockdown and MAPK inhibition, neither mesodermal nor endodermal markers were expressed. Instead, zygotic markers of the aboral ectoderm *six3/6* and *foxQ2a* were expressed ubiquitously (Figures 4E and 4F). This is striking since earlier studies showed that aboral ectoderm markers are abolished in the β-catenin morphants ^15,33^. Moreover, this ubiquitous expression of *six3/6* when both MAPK and β-catenin signaling are inhibited, appears similar to what happens when aboral cells, isolated by bisection at the eight-cell stage, growth in the absence of signals from the oral domain ^15^. Our new results suggest that i) MAPK-dependent suppression of *six3/6* and *foxQ2a* aborally is caused by the expansion of the mesoderm in the β-catenin morphants; ii) show that aboral ectoderm represents the default state of the embryo in the absence of MAPK and β- catenin signalings; and iii) indicate that MAPK signaling activity is what activates mesodermal genes in the β-catenin morphants (Figure 4F).

### Endoderm segregation involves the Notch pathway

In *Nematostella*, the endoderm emerges between the mesoderm and ectoderm around 12-14 hpf. As the Notch pathway is known to play a role in segregation of mesoderm and endoderm in other organisms ^38^, we investigated its function in endoderm initiation in *Nematostella*. First, we conducted an expression analysis of the receptor *notch* and its potential ligands at multiple time points from egg to 26 hpf using in situ hybridization. While only one gene coding for the Notch receptor has been previously found in *Nematostella* ^39^, four ligands of this pathway, *delta*, delta-like, *jagged1-like* and *jagged1B-like*, can be identified in the genome. In situ hybridization shows that *notch* mRNA is maternally deposited in the egg and persists in the embryo. It is initially expressed ubiquitously, but then its expression disappears from the mesoderm around 10hpf (Figure 5A and S5A). After gastrulation, Notch expression is downregulated in the midbody ectoderm ^39^ (Figure S5). mRNAs coding for *delta* and *delta-like* are zygotic, but they are also expressed before the onset of gastrulation, at 10 and 14 hpf, respectively. Both are primarily found in the mesoderm, with *delta-like* mRNA transcripts visualized from 10hpf (Figure 5A and S5A). In addition, *delta* (but not *delta-like*) is also detected at the aboral side in a salt and pepper pattern after 12hpf, reflecting its role in early neurogenesis in this domain ^39,40^. The early mesodermal expression of *delta* shows a peak between 14-16hpf before its expression gradually decreased until gastrulation (Figure 5A and S5A).The third Notch ligand gene, *jagged1-like*, exhibits a later, and rather weak expression starting at 18hpf in the mesoderm (Figure S5A). While *delta* and *jagged1-like* expression decreased in the mesoderm after 18hpf, *delta-like* expression starts at 16 hpf and continues to be abundantly detected in the mesoderm after gastrulation (Figure S5A). Since the last Notch ligand gene *jagged1B-like*, based on single cell RNA sequencing data, is expressed at very low level uniformly across all cell clusters, we did not investigate its expression at early stages. Thus, notch ligands *delta, delta-like* and *jagged1-like* are mostly expressed in the mesoderm, whereas *notch* is expressed in the ectoderm. Double in situ hybridization shows that the expression of *delta* in the mesoderm and *notch* in the whole ectoderm of the embryo are precisely complementary, without any overlap (Figure 5B). To assess, whether endoderm is induced on the mesodermal or the ectodermal side of the *notch-delta* boundary, we analyzed the expression of *notch* together with the endodermal marker *foxA*. We detect *foxA* expression in the *notch*-expressing ectodermal border cells (Figure 5C), which are in direct contact with the *delta*-expressing mesodermal cells (Figure 5B). These results suggest that endoderm is ectoderm-derived, and that Notch signaling may have a role in endoderm induction.

**Figure 5.**
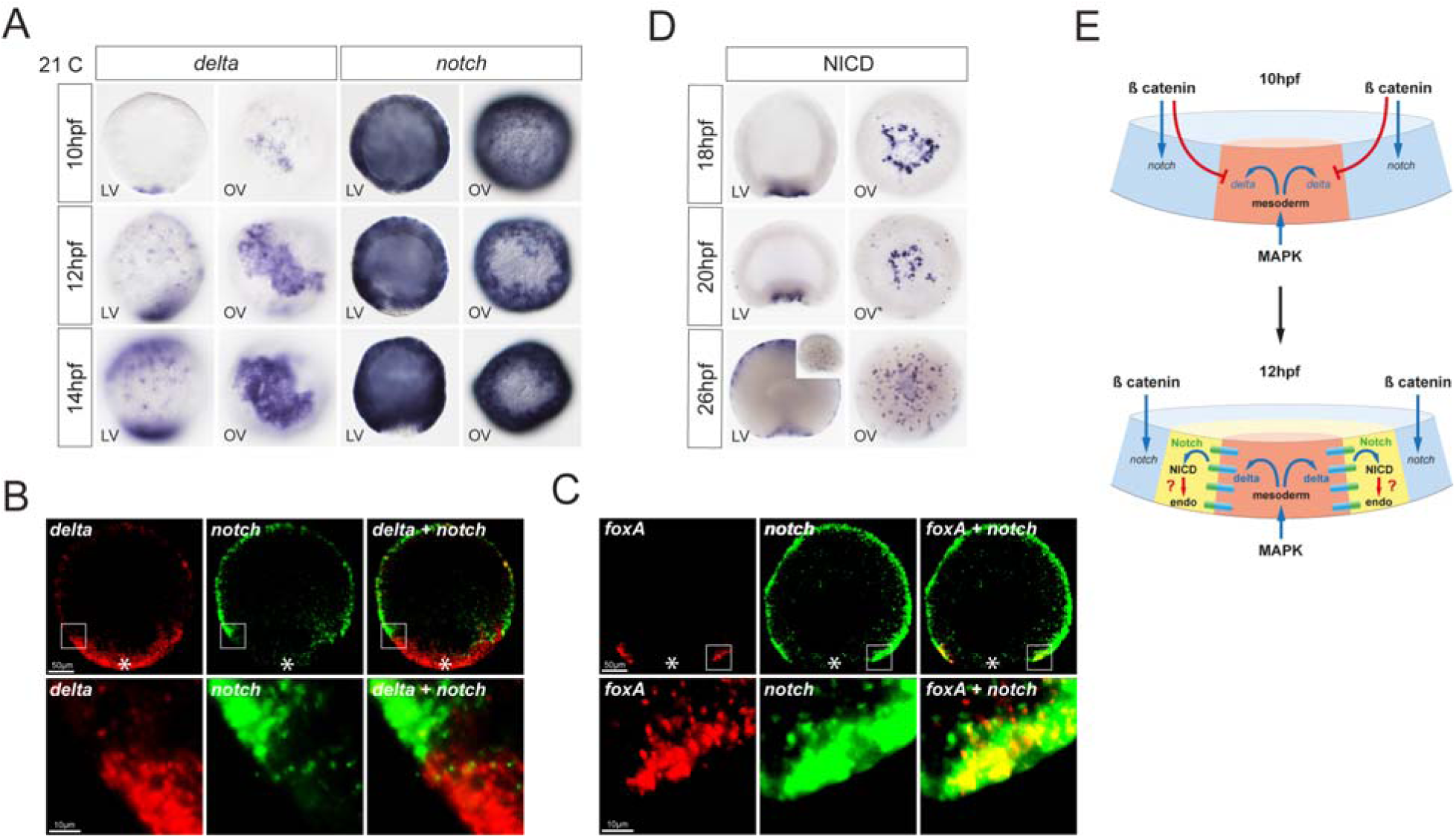
Delta and Notch expression demarcate the mesoderm/ectoderm border. (A) Expression profile analysis of the ligand *delta* and receptor (*notch*) of the Notch pathway from 10hpf to 14hpf by in situ hybridization. While *delta* (and *delta-like)* are expressed mostly in the mesoderm, *notch* is detected in the ectoderm. LV; Lateral view, OV; oral view. (B) Double in situ fluorescence of *delta* (red) and *notch* (green) expression. *delta* (mesoderm) and *notch* (ectoderm) are expressed in a complementary, abutting pattern. (C) Double in situ fluorescence of *foxA* and *notch. foxA* is expressed in the ectodermal border cells, which are in contact with the mesoderm and expressing *notch*. (D) Activated Notch pathway is revealed with NICD immunostaining at 18, 20 and 26hpf to. LV; lateral view, OV; oral view. (E) Schematic summary of the regulation of *delta* and *notch* expression by β-catenin and MAPK signaling following by endodermal genes activation by the Notch pathway.

The activated Notch pathway is marked by the translocation of the Notch intracellular domain (NICD) to the nucleus (Figure 5D). To check, which cells and domains in the embryo show an activated Notch pathway, we carried out immunohistochemistry against the NICD in early embryos. Strikingly, we find two domains in early gastrula stage embryos: first, an uninterrupted ring at the blastopore corresponding to the endoderm/mesoderm *Notch* expression border at late blastula stage (Figure 5D) and second, several hours later, small cell patches in the aboral half, consistent with the second role of Notch signaling in specifying neuronal precursor cells (Figure 5D; ^39,41,40^. To gain a better understanding of the involvement of the Notch signaling pathway in endoderm formation, we examined its regulation by the Wnt/β- catenin signaling pathway. Upon Morpholino mediated knockdown of β*-catenin, delta* and *delta-like* expression became ubiquitous, while *notch* expression was lost (Figure S5B). Conversely, overactivation of β-catenin signaling by AZ treatment results in the loss of *delta* and *delta-like* and the extension of *notch* expression throughout the whole embryo. Thus, *delta* and *delta-like*, like other mesodermal genes, are repressed, while *notch*, like other earlier ectoderm genes, is activated by the β-catenin signaling (Figure 5E and S5B). By contrast, a knockdown of the MAPK signaling pathway target and important mesodermally expressed transcription factor *erg*, did not lead to significant changes in the expression of the *delta* gene. However *erg* knockdown induces ectopic expression of *notch* in the mesoderm, suggesting that although ERG does not contribute significantly to the regulation of *delta* expression it may still be involved in the repression of *notch* in the mesoderm (Figure S5B).

### Endoderm is induced by Delta-Notch signaling between ectoderm and mesoderm

The formation of the endoderm on the *notch* side of the *delta*-*notch* expression boundary raises the possibility that the activation of the Notch pathway by Delta induces endoderm formation. To test this, we inhibited the Notch pathway with γ-secretase inhibitors LY-411575 and DAPT, which prevent the release of the intracellular domain of the Notch receptor (NICD) (Figure 6A). Upon treatment with increasing concentrations of Notch signaling inhibitors, the expression of endodermal genes *brachyury*, *foxA*, *wnt1* and *wnt3* at 20hpf was repressed in a concentration-dependent manner (Figure 6B). By contrast, Notch signaling inhibition had no effect on the expression of the mesodermal genes, such as *snailA* or *zicA* (Figure 6B). In line with the previous observations, the disappearance of the *wnt*- and *brachyury*- expressing endodermal fate did not affect gastrulation but led to defects in the Wnt-dependent patterning of the ectoderm and affected pharynx formation: we observed the shift of the midbody marker *Wnt2* expression towards the oral end and the expansion of the aboral ectoderm markers *foxQ2a* and *Six3/6* at the gastrula stage, and suppression of the endodermal pharynx development at the 48 hpf planula stage (Figures 6B-C and Figure S6; ^15,24,42–44)^. Thus, we conclude that Notch signaling is necessary for endodermal fate induction in the ectodermal cells located at the border to the mesoderm.

**Figure 6.**
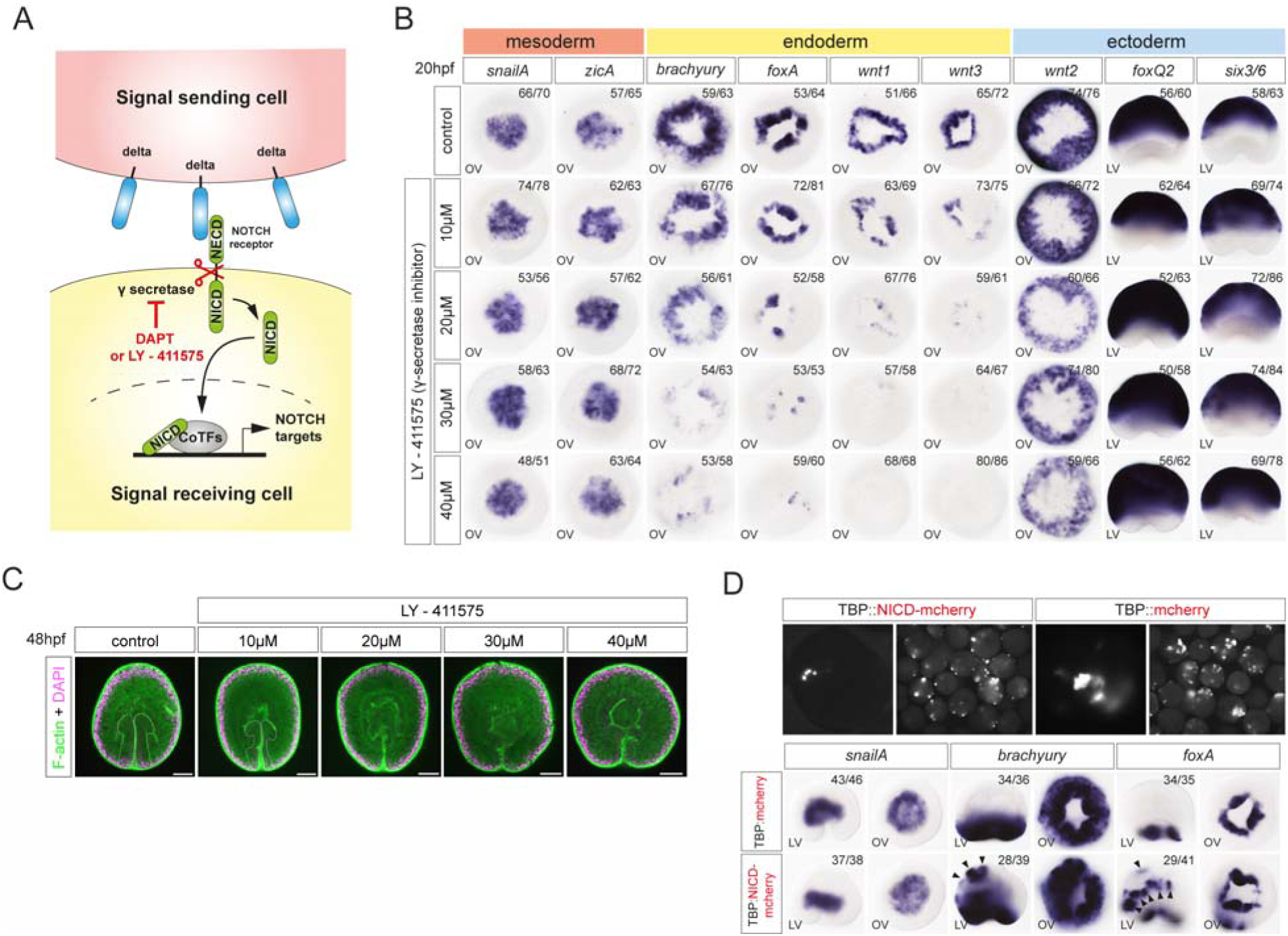
Notch signaling pathway plays a crucial role in endodermal tissue formation by activating endodermal genes. (A) Scheme of Notch signaling and the target of the inhibitor LY-411575. (B) Visualization of mesodermal (*snailA, zicA*), endodermal (*brachyury, foxA, wnt1, wnt3, wnt1*) and ectodermal (*wnt2, foxQ2, six3/6*) gene expression at 20hpf on embryos treated with LY-411575 using in situ hybridization; Expression of endodermal genes is downregulated dose-dependently upon inhibition of the Notch signaling. LV; lateral view, OV; oral view. (C) Morphological analysis on LY-411575 treated embryos stained with phalloidin (green) and DAPI (cyan). LY-411575 treatment induce the loss of the endoderm; Scale bar, 100 μm. (D) Analysis of mesodermal (*snailA*) and endodermal (*brachyury, foxA*) gene expression in *TBP::mCherry* and *TBP::NICD-mCherry* injected embryos at 20hpf. Black arrow heads show ectopic expression of endodermal markers when NICD is overexpressed. LV; lateral view, OV; oral view.

To test whether Notch signaling is also sufficient to induce endoderm, we ectopically activated Notch signaling in the ectoderm by mosaically overexpressing either *NICD-mCherry* or, as a negative control, just *mCherry* under control of the ubiquitously active *TBP* promoter using the established transgenesis protocol ^25,45^. The visualization of mCherry expression showed a uniform distribution throughout the cells, whereas NICD-mCherry clearly suggested nuclear translocation. Strikingly, injection of *TBP::NICD-mCherry* plasmid resulted in ectopic expression of *foxA* and *brachyury* in 60% of cases, whereas no ectopic endodermal marker expression was observed with the *TBP:: mCherry* plasmid (Figure 6D). The expression of *snailA* was not affected, indicating that NICD specifically activates endodermal and not mesodermal genes (Figure 6D). Taken together, these experiments show that Notch signaling is both necessary and sufficient to induce endodermal genes in the Notch-expressing ectodermal domain, consistent with the idea that the abutting expression of mesodermal *delta* and ectodermal *notch* at 10-14 hpf leads to the induction of endoderm.

## Discussion

In this study, we investigated how three signaling pathways – β-catenin, MAPK, and Notch – interact and function during the specification of the three germ layer identities in *Nematostella*. We show that in *Nematostella*, mesoderm is the first embryonic territory to be specified in the 6 hpf blastula stage embryo, which has ubiquitous aboral ectoderm identity as a default state. While the activation of the mesoderm specification program in a very precisely defined β- catenin-negative domain appears to rely on yet unknown maternal components ^16^, the role of β- catenin signaling in the early blastula is to prevent mesodermal gene expression outside of the mesodermal domain (^15^; this study) and, as we show here, to promote expression of the early ectodermal markers outside the mesoderm. Using pharmacological treatments and gene knockdown, we demonstrate that MAPK signaling is responsible not only for the activation of the mesodermal marker expression but also for the repression of the zygotically expressed ectodermal markers such as *six3/6* and *foxQ2a*. Finally, we show that endodermal fate is induced in a single row of cells at the interface between the Notch-expressing ectoderm and the Delta-expressing mesoderm. Moreover, Notch signaling activation is necessary and sufficient for the initiation of the endodermal gene expression program in any ectodermal cell. We consider the latter two findings highly significant and will discuss them below.

### MEK/MAPkinase signaling, an ancestral and conserved pathway to induce mesoderm

We showed that activation of the MAP kinase signaling is essential for the specification of the mesoderm in the cnidarian *Nematostella*. Its inhibition with a selective inhibitor of MEK disrupts early mesodermal gene expression and causes gastrulation defects. We see a similar situation in some Bilateria, especially in ambulacrarian deuterostomes (Hemichordata and Echinodermata), whose germ layer fate maps – mesodermal cells at the vegetal pole surrounded by a concentric endodermal domain, and an ectoderm making up the rest of the embryo beyond the endoderm – are strikingly similar to the germ layer fate map of *Nematostella* (Figure 7), except that in *Nematostella* meso- and endoderm form at the animal rather than at the vegetal pole ^46–49^, MAPK signaling was shown to be important for the formation of the primary and of a subset of the secondary mesenchymal cells in sea urchin *Paracentrotus* ^50^, and for the mesoderm specification in the hemichordates *Saccoglossus* and *Ptychodera* ^51–53^. The situation in non-vertebrate chordates is somewhat different. FGF-dependent MAPK signaling is responsible for the mesoderm induction by the future endodermal cells in the urochordate *Ciona* ^54,55^, while in the cephalochordate *Branchiostoma* only anterior somitic mesoderm appears to form in a MAPK-dependent manner ^56^. In vertebrates, Nodal and TGFß signaling are responsible for mesoderm induction, however, FGF-mediated MAPK signaling (together with Wnt, BMP and Hedgehog signaling pathways) is involved in the mesodermal patterning ^57,58^. In protostomes, the role of FGF-mediated MAPK signaling has been investigated in three lophophorate spiralian species: the brachiopods *Terebratalia* and *Novocrania* and in a related phoronid *Phoronopsis*. In the two brachiopods, pharmacological suppression of the FGF receptor resulted in embryos lacking coelomic mesoderm, and muscle tissue was also lost in the phoronid embryo ^13^. In another spiralian, the annelid *Alitta*, MAPK inhibition led to defects in the mesodermal patterning and morphogenesis, however, it did not abolish the mesodermal fate ^59^. MAPK signaling is well-known for its role in the specification of the 3D macromere in gastropods, such as the snails *Crepidula* and *Ilyanassa* ^60,61^ . However, the daughter cell of the 3D, the mesentoblast 4d, appears not to require MAPK signaling, and although heart and muscle are missing in *Ilyanassa* upon MEK inhibitor treatment, we cannot be certain that the role of MAPK signaling here is induction of mesoderm. Nevertheless, the role of MAPK signaling in mesoderm specification has been documented in ambulacrarian deutorostomes, in lophophorate spiralian protostomes and now also in a cnidarian *Nematostella*. Thus, despite all the diversity, it appears plausible that the origin of MAPK-signaling as a mechanism of specification of mesoderm tissue identity may have predated the split of Cnidaria and Bilateria.

**Figure 7:**
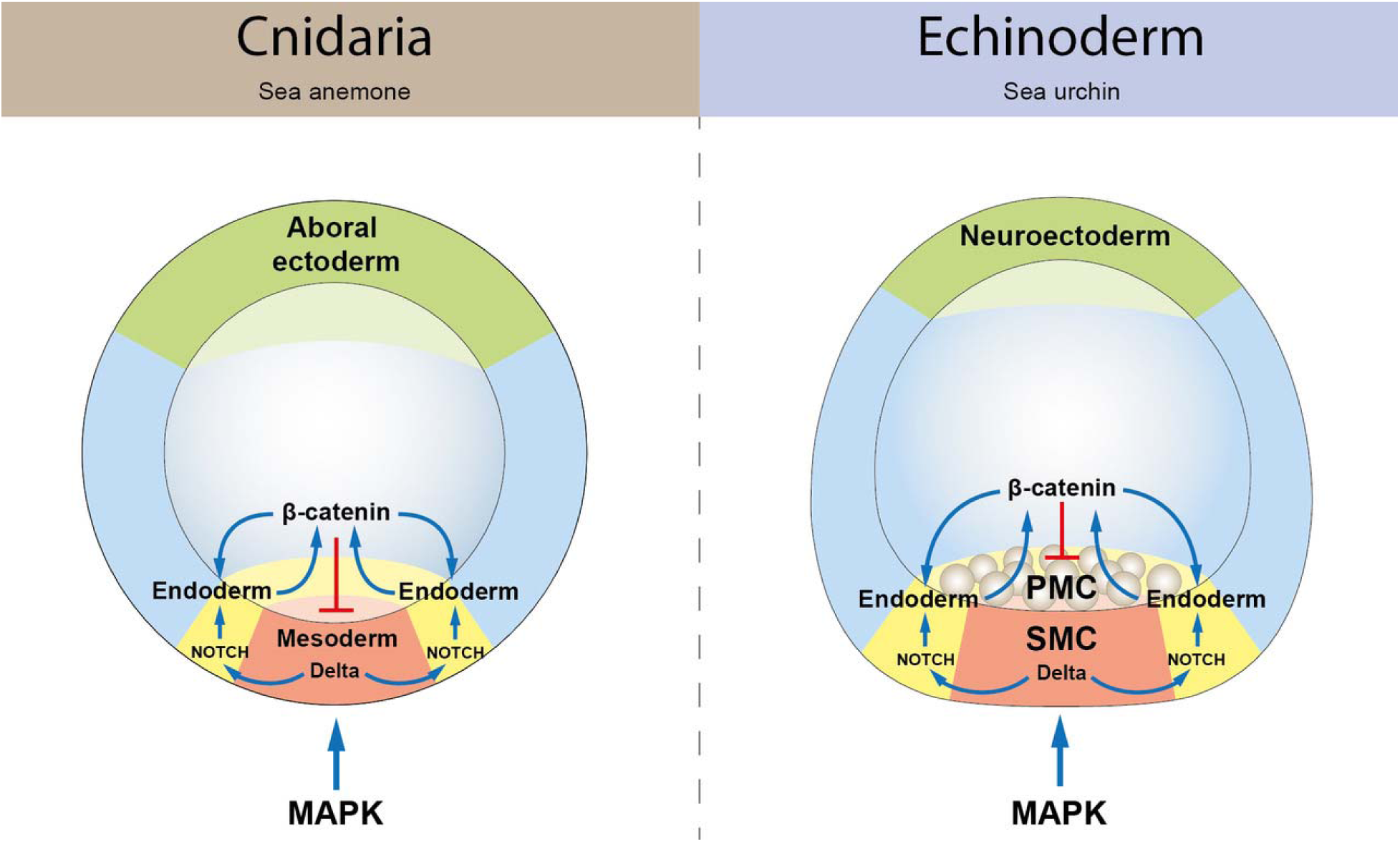
Comparison of interactions of signaling pathways in the specification of the sea anemone *Nematostella* and sea urchins.

### Notch signaling induces endoderm specification in *Nematostella*

As mentioned above, in terms of mutual positions of the germ layers, the fate map of the early *Nematostella* embryo is strikingly similar to the fate maps of ambulacrarian deuterostomes. Despite the difference in the mechanism of the mesoderm specification (β- catenin dependent endomesoderm specification followed by turning β-catenin signaling off in the mesoderm once it is specified and keeping β-catenin signaling on in the endoderm in Ambulacraria vs. mesoderm specification in a domain initially lacking nuclear β-catenin in *Nematostella*), the subsequent patterning of the endoderm and ectoderm by a gradient of Wnt/β-catenin signaling appears to use the same logic and activate orthologous downstream genes ^16,24^. However, this raises an obvious question: when and how does *Nematostella* forms the endoderm if mesoderm is specified in the β-catenin-negative domain of the early blastula, and the remaining maternal β-catenin-positive domain represents the ectoderm? Our data show that this occurs several hours later by induction.

One striking feature shared by ambulacrarian and cnidarian embryos is that pharmacological upregulation of β-catenin signaling leads to endodermal marker expression in the ectodermal cells, while mesodermal cells, once specified, become insensitive to changes in the β-catenin levels. The best candidate signaling pathway for inducing a novel fate at the interface of two different populations of cells was the Delta/Notch pathway. In animal development, Delta/Notch signaling is a major facilitator of binary cell fate decisions using two distinct types of regulatory logic: lateral inhibition and lateral induction. During lateral inhibition, a cell within an “equivalence group” of equipotent cells expressing low levels of Notch and Delta is induced to acquire a certain fate, which makes it produce more Delta than its neighbors. This Delta-producing cell signals to the neighbors causing them to shut down Delta production and acquire an alternative fate ^38^. This mechanism is often used in neural differentiation, likely including its previously reported role in the formation of the nervous system in *Nematostella* ^39–41^. During lateral induction, scenarios may be different. In a field of equipotent cells expressing both Delta and Notch, lateral induction may result in concerted behavior of large groups of cells as in vertebrate somitogenesis and, possibly, in arthropod segmentation ^62^. However, at the interface between the Notch-expressing cell population and the Delta-expressing cell population, lateral induction leads to the emergence of the third cell state ^63^ – in *Nematostella*, this is the endoderm. In sea urchins, like in *Nematostella*, Notch is initially ubiquitous, and eventually suppressed in the mesodermal lineage, where Delta starts to be expressed, however, unlike in *Nematostella*, Delta/Notch signaling is responsible for the segregation of the specific mesodermal populations rather than for the induction of the endodermal fate ^9,38,64,65^. In contrast, in sea stars, which lack the skeletogenic primary mesenchymal cells, Delta/Notch signaling is responsible for the formation of the endoderm/mesoderm boundary ^11^ despite a similar expression pattern of *Notch* and *Delta* as in sea urchins. Similarly, Delta signaling from mesoderm to endoderm and Notch-dependent suppression of the mesodermal gene expression in the endoderm has been hypothesized for cephalochordates ^66^. Thus, whether Notch-dependent endoderm/mesoderm segregation represents a trait ancestral for Cnidaria and Bilateria remains difficult to judge. Delta/Notch signaling is an efficient boundary formation mechanism, which can be used differently in the process of endoderm-mesoderm segregation even in the closely related animal clades. However, mesodermal expression of *Delta* versus endo- and ectodermal expression of *notch* may indeed represent an ancestral condition.

### Comparison of germ layer identities in *Nematostella* and bilaterians

An earlier study has challenged long held views about homologies of cnidarian and bilaterian germ layers and postulated that the inner cell layer, often termed endoderm (or gastrodermis) in the cnidarian literature, has a molecular profile very reminiscent of bilaterian mesoderm^1^. In line with that, the cellular behavior (partial or full EMT) during gastrulation as well as its cellular derivatives (gonads, muscles, storage tissue) is characteristic of mesoderm. Our study lends further support for the previously postulated segregation of three germ layer identities in a diploblastic animal. We show that also its early activation by MAPK is shared with mesoderm induction in many bilaterians. We therefore propose to term this tissue mesoderm in *Nematostella*.

Of note, in many bilaterians, there is a mesendodermal state during early development, which then segregates into distinct endoderm and mesoderm ^67^, while in *Nematostella*, we only observe a very weak and transient expression of a few wnt genes in the future mesoderm, before they resume in the ring of endodermal tissue surrounding the mesoderm. However, nuclear β-catenin is barely detectable in this domain ^16^, suggesting that the short and low level expression of a few wnt mRNAs does not lead to activation of the Wnt/β-catenin pathway. We suppose that MAPK signaling suppresses further strengthening of wnt gene expression before they can even start signaling through the wnt pathway. Thus, it is questionable whether there is a true mesendodermal state in *Nematostella*. By contrast, the endodermal tissue identity is induced in the ectodermal side of the boundary to the mesoderm. This is in line with our single cell RNAseq data, which show a stronger link of endodermal identity with the rest of the ectoderm, while mesoderm is already very distinct. Moreover, this tissue and its derivative, the septal filament, also produces cnidocytes, which typically only occur in ectoderm. Thus, the endodermal cell fate appears to retain some features typical of ectoderm. While the lack of mesendodermal state might appear as a difference to many bilaterians, there are also examples, in particular among the ecdysozoans, where fore- and hindgut derive from ectodermal origin ^69,70^ and continue to display some ectodermal features, such as chitinization of the epithelial layer. Thus, despite its diploblastic nature, *Nematostella* uses conserved signaling mechanisms to segregate three germ layers as many bilaterians. Preliminary evidence suggests that this is also shared in members of the other major branch of cnidarians, the scyphozoans ^1^, but further studies are required to confirm this.

## Conclusion

Taken together, we find that during germ layer specification in the sea anemone *Nematostella vectensis* MAPK signaling in the β-catenin negative background specifies the mesodermal and represses the ectodermal fate, while Notch signaling induces endodermal fate in the ectodermal cells bordering the mesoderm. We propose that involvement of MAPK signaling in mesoderm formation represents an ancestral trait shared by Cnidaria and Bilateria. In contrast, although Notch and Delta are recurrently involved in endo- and mesoderm formation in different models, a wider phylogenetic sampling is required to decide whether Delta/Notch dependent endoderm specification is ancestral for Cnidaria+Bilateria.

## Acknowledgements

This research was funded in whole or in part by the Austrian Science Fund (FWF) grants to U.T. (P34404) and Austrian Science Foundation (FWF) grants 10.55776/P30404 and 10.55776/P36080 to G.G.. T.L. was a recipient of the fellowship ICM-2017-07957 of the Stipendienstiftung der Republik Österreich. For the purpose of Open Access, the authors have applied a CC BY public copyright license to any Author Accepted Manuscript (AAM) version arising from this submission.

---

**Figure S1.**
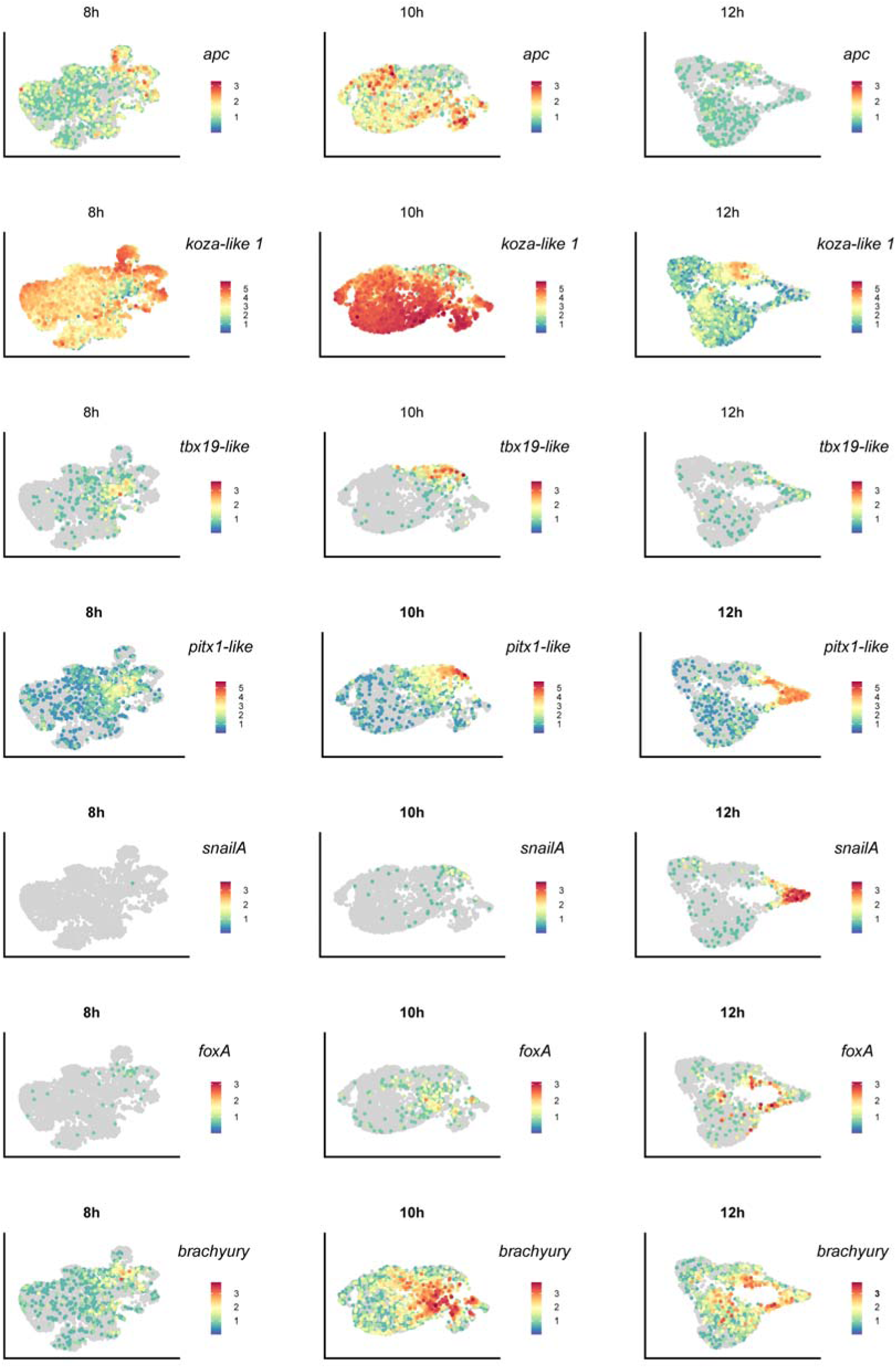
UMAP projections of single RNAseq datasets at 8, 10 and 12hpf. Visualization of the cells expressing ectodermal genes (*apc, koza-like 1*), mesodermal genes (*tbx19-like, pitx1-like, snailA*) and endodermal genes (*foxA, brachyury*) at different time points during early development in *Nematostella*.

**Figure S2.**
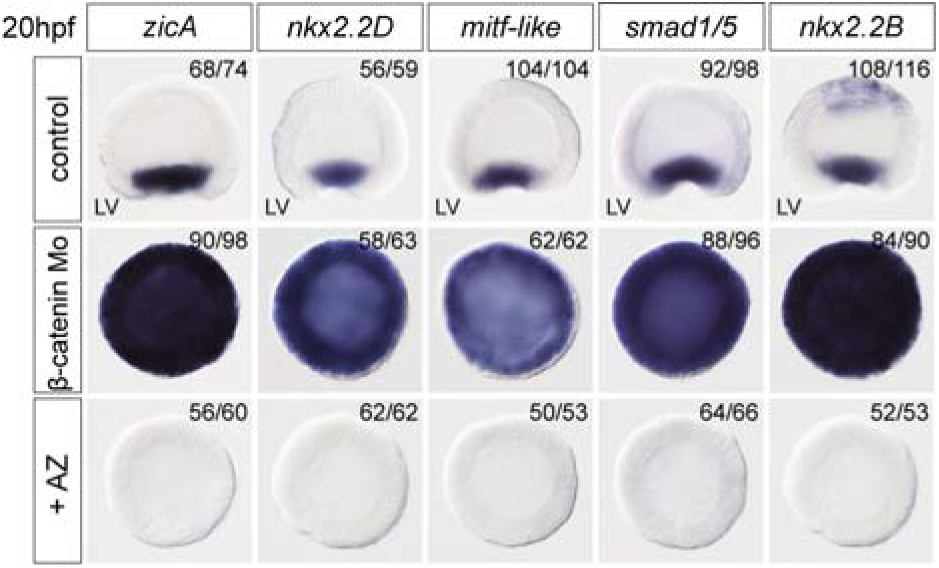
β-catenin signaling prevents mesodermal marker expression at 20 hpf. Expression of mesodermal (*zicA*, *nkx2.2D*, *mitf-like*, *smad1/5*, *nkx2.2B*) genes in AZ treated embryos or β-catenin morpholino embryos at 20hpf. LV; lateral view.

**Figure S4.**
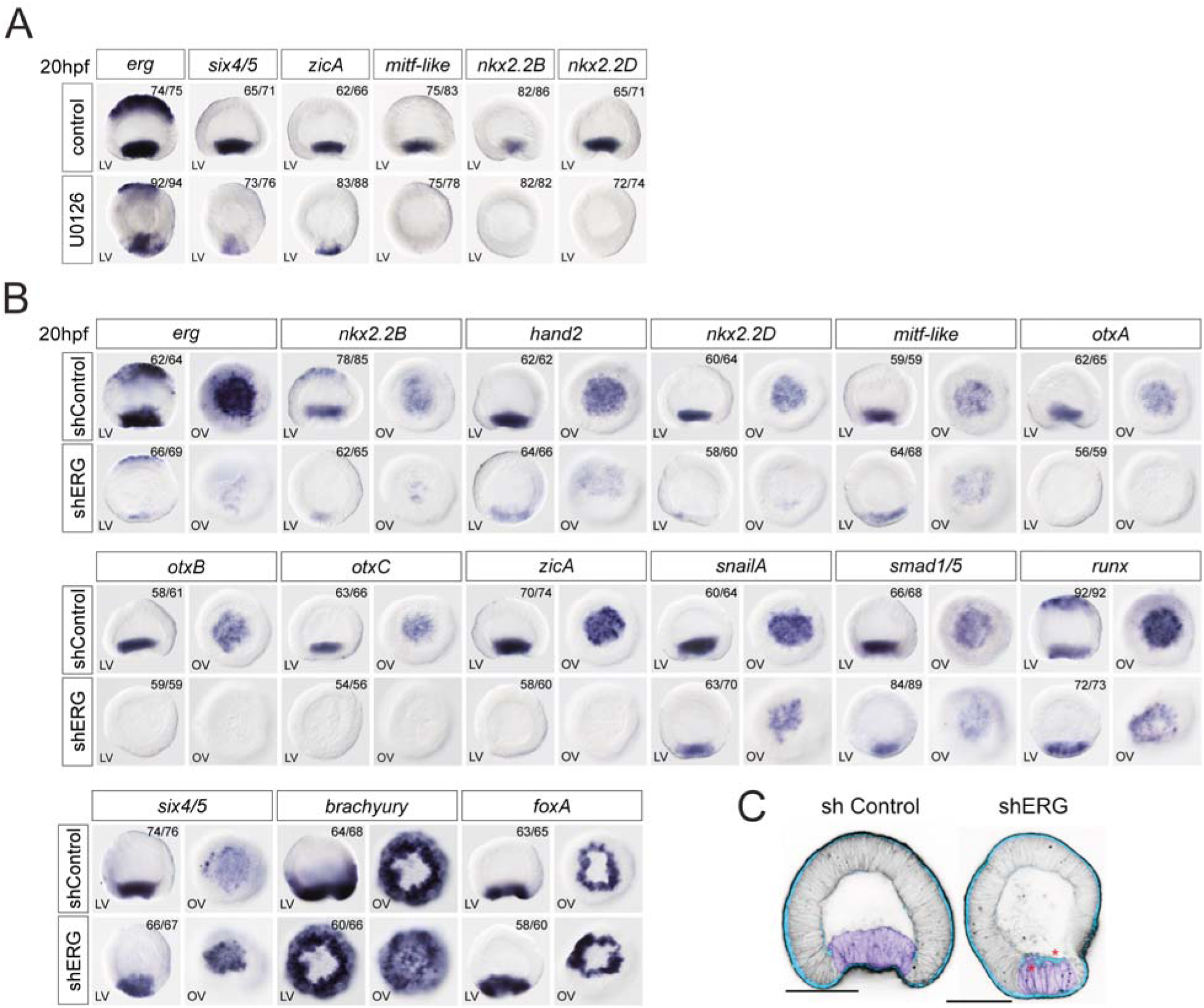
ERG is essential for mesoderm specification and morphogenesis. (A) Mesodermal (*erg*, *six4/5*, *zicA*, *mitf-like*, *nkx2.2B*, *nk2.2D*) gene expression analysis at 20hpf with in situ hybridization on U0126 treated embryos. LV; lateral view. (B) Mesodermal (*erg*, *nkx2.2B*, *hand2*, *nkx2.2D*, *mitf-like*, *otxA*, *otxB*, *otxC*, *zicA, snailA, smad1/5, runx*, *six4/5*) and endodermal (*brachyury*, *foxA*) gene expression analysis at 20hpf with in situ hybridization on shERG injected embryos. LV; lateral view, OV; oral view. (C) The effect of the ERG knockdown on cdh3 expression (blue) and on gastrulation movements phenocopies the effect of the MAPK signaling inhibition by U0126. F-actin (black) staining is used to show the cell outlines.

**Figure S5.**
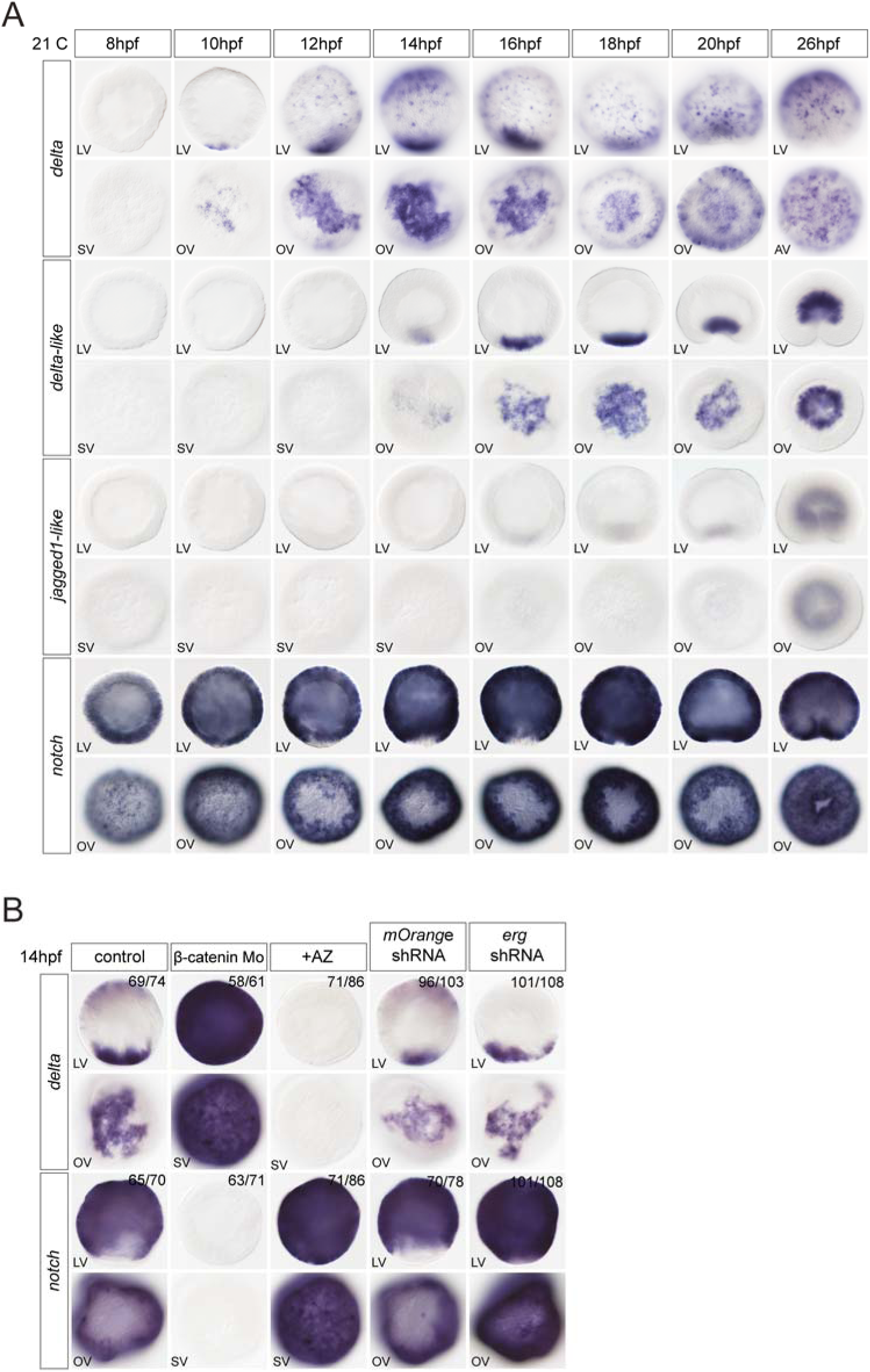
Regulation of ligands and receptor of the Notch pathway. (A) Expression analysis of genes coding for ligands (*delta, delta-like, jagged1-like*)and receptor (*notch*) of the Notch pathway from 8hpf to 26hpf by in situ hybridization. LV; lateral view, OV; oral view. (B) In situ hybridization at 14hpf in AZ treated embryos, β-catenin morpholino or shRNA against *erg* injected embryos to visualize mesodermal (*delta*) and ectodermal (*notch*) genes expression. LV; lateral view, OV; oral view, SV; surface view.

**Figure S6.**
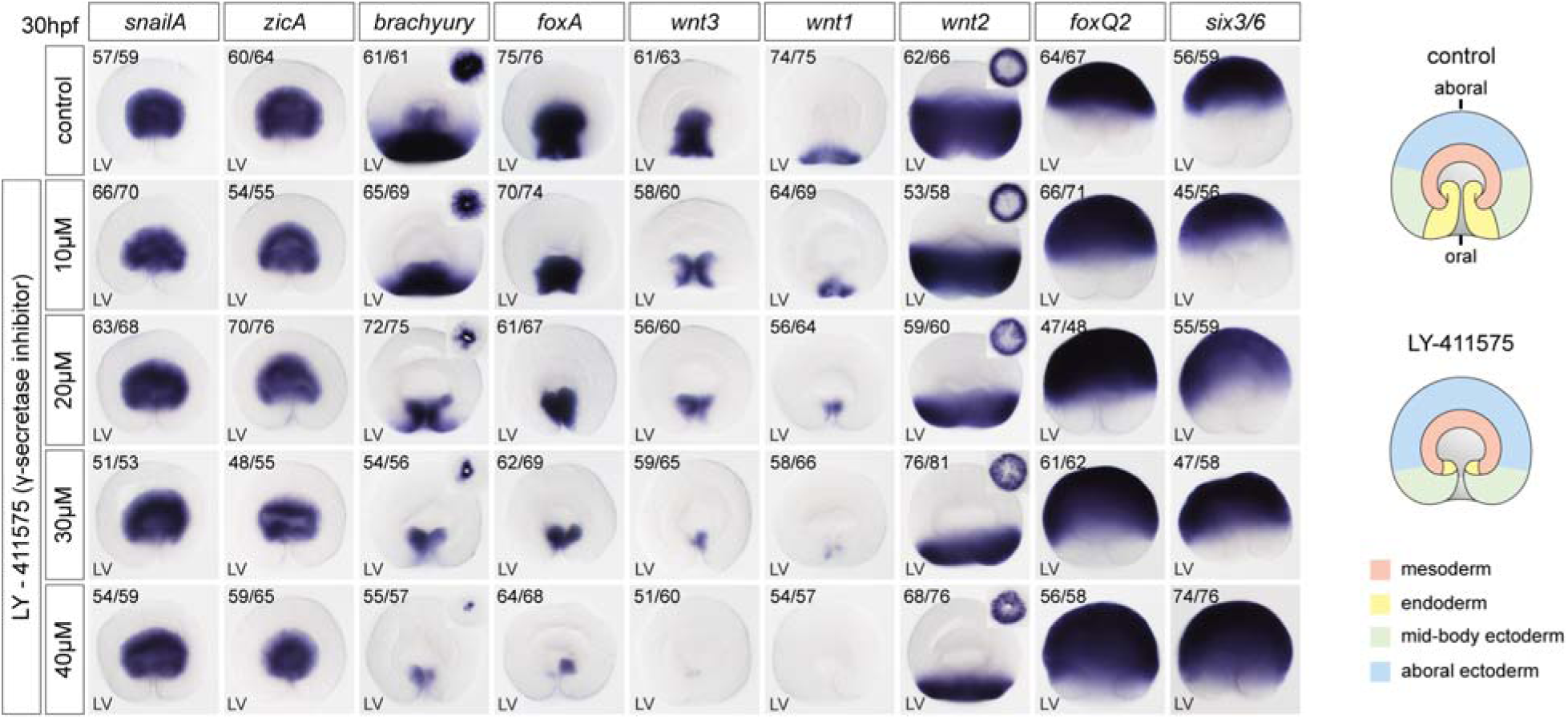
Notch signaling is essential for endodermal gene expression. Mesodermal (*snailA, zicA*), endodermal (*brachyury, foxA, wnt1, wnt3, wnt1*) and ectodermal (*wnt2, foxQ2, six3/6*) gene expression analysis at 30hpf on embryos treated with LY-411575 using in situ hybridization; Expression of endodermal genes is downregulated after inhibition of the Notch signaling. LV; lateral view, OV; oral view.

